# Molecular Basis for the Interaction of Catalase with D-Penicillamine : Rationalization of some of its Deleterious Effects

**DOI:** 10.1101/2021.09.16.460603

**Authors:** Dominique Padovani, Erwan Galardon

## Abstract

D-penicillamine (D-Pen) is a sulfur compound used in the management of rheumatoid arthritis, Wilson’s disease (WD), and alcohol dependence. Many side effects are associated with its use, particularly after long-term treatment. However, the molecular bases for such side effects are poorly understood. Based on the well-known oxidase activity of hemoproteins, and the participation of catalase in cellular H_2_O_2_ redox signaling, we posit that D-Pen could inactivate catalase, thus disturbing H_2_O_2_ levels. Herein, we report on the molecular bases that could partly explain the side effects associated with this drug compound, and we demonstrate that it induces the formation of compound II, a temporarily inactive state of the enzyme, through two distinct mechanisms. Initially, D-Pen reacts with native catalase and/or iron metal ions, used to mimic non heme iron overload observed in long-term treated WD patients, to generate thiyl radicals. These partake into a futile redox cycling, thus producing superoxide radical anions O_2_^•-^ and hydrogen peroxide H_2_O_2_. Then, either H_2_O_2_ unexpectedly reacts with native CAT-Fe(II) to produce compound II, or both aforementioned reactive oxygen species intervene into compound II generation through compound I formation then reduction. These findings support evidence that D-Pen could perturb H_2_O_2_ redox homeostasis through transient but recurring catalase inactivation, which may in part rationalize some deleterious effects observed with this therapeutic agent, as discussed.

**Graphical abstract:** 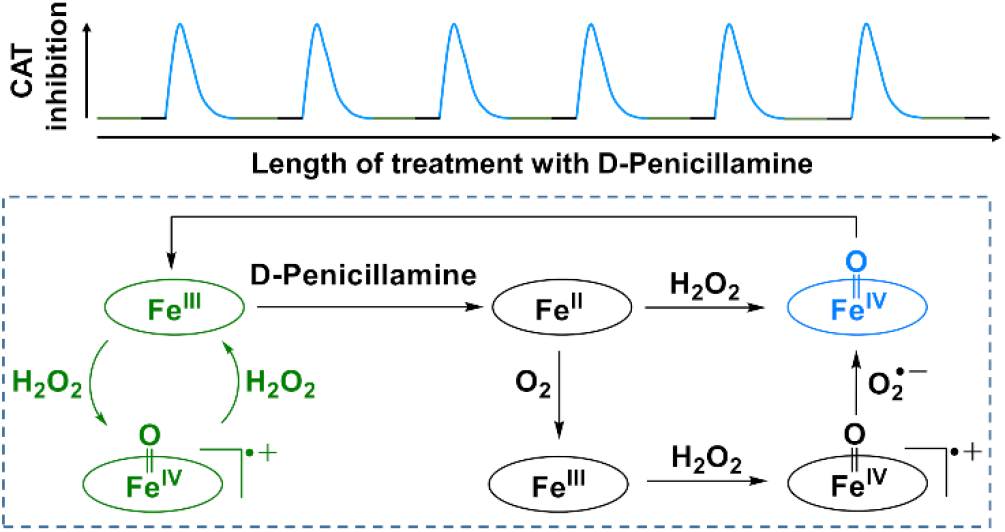

## Introduction

D-penicillamine (D-Pen) is an unnatural thiol mostly implicated in three types of reactions when used in a biological context: thiol-disulfide exchange, thiazolidine formation in the presence of carbonyl compounds, and metal chelation (**1**,**2**). As a result, D-Pen is used as a therapeutic copper chelating agent in the treatment of Wilson’s disease (WD), an autosomal recessive inherited copper metabolism disorder that results in severe defects such as oxidative tissue damage, liver damage (chronic hepatitis and cirrhosis), and ultimately death if untreated (**3**).

D-Pen is also used as an immunosuppressor in the treatment of rheumatoid arthritis (RA), a chronic inflammatory disease of the joints in which reactive oxygen species (ROS) as well as various enzymes released at the inflamed joints (*e.g*. heme-dependent myeloperoxidase, MPO) may contribute to the etiology of the disease (**4**-**5**). D-pen may exert a modulatory function on the oxidant-producing capacity of MPO by scavenging MPO-generated hypochlorous acid and by reacting with redox intermediates, thus contributing to superoxide radical anions (O_2_^•-^) production and then inactive compound III (MPO-Fe(III)-O_2_^•-^) generation (**6**).

Last, D-Pen has been reported to act as a sequestration agent of acetaldehyde (ACD), a metabolite of ethanol generated either through the action of alcohol dehydrogenase in the liver or *via* the peroxidatic activity of catalase (CAT) in the brain (**7**-**8**). Brain derived ACD acts as a neuroactive agent in the central nervous system (**9**). As a result, strategies have been established for blocking ethanol metabolic pathways (e.g. inhibition of catalase bioactivity) and trapping ACD (*e.g*. sequestering of ACD by D-Pen through thiazolidine formation) (**8**-**9**).

Unfortunately, although the D-enantiomer does not display the acute toxicity encountered with the use of the L-enantiomer, a broad spectrum of toxicities has been reported for D-Pen particularly with increasing time of treatment (**2**-**4**). Thus, D-Pen exhibits severe secondary effects such as hepatotoxicity (**10**), immune disorders (**11**), and high incidence of neurological worsening (**12**-**13**). Additionally, hepatic non-heme iron overload occurs in patients presenting with WD during long-term treatment with D-Pen (**14**). This combination of copper and iron perturbed homeostasis may play a significant role in the pathogenesis of acute and chronic liver damage observed in animal models of WD (**15**). Also, a recent study suggests that D-Pen could selectively perturb hydrogen sulfide homeostasis, thus reducing vasodilation in mouse artoa rings, or exacerbating the inflammatory response in a mouse model of inflammation (**16**).

Noteworthy, some of D-Pen side effects could also be explicated by considering a D-Pen-induced alteration of hydrogen peroxide (H_2_O_2_) homeostasis. Indeed, while H_2_O_2_ is a well-recognized signaling messenger (**17**), it is also well-known for its cytotoxic effects and its contribution to the etiology of immune diseases, its involvement in vasoconstriction, and its participation to a time-dependent increase in the cellular labile iron pool in neuronal cells (**18**-**20**). On the basis of the well-known thiol oxidase activity of hemoproteins (**21**), including catalase (**22**), and the participation of catalase in cellular H_2_O_2_ redox signaling (**23**), we posit that D-Pen could induce catalase inactivation and thus disturb H_2_O_2_ homeostasis, which could play a key role in the reported cytotoxicity of D-Pen. Accordingly, we deciphered the reactivity of D-Pen with catalase.

We report here that catalase is indeed inactivated by D-Pen with a pharmacologically relevant relative IC_50_ value. The reactivity of catalase with D-Pen first involves D-Pen oxidation to its corresponding thiyl radical. The latter next enter a futile redox cycling that further facilitates the formation of compound II (CAT-Fe(IV)=O), a temporarily inactive state of catalase. Interestingly, the presence of iron metal ions, used to mimic hepatic non-heme iron overload occurring in WD patients treated with D-Pen, has a significant effect on the inhibitory capacity of D-Pen, leading to a drastic drop in the relative IC_50_ value. Surprisingly, the generation of compound II occurs through two distinct mechanisms, one of which is unusual. Our findings support evidence that D-Pen could momentarily perturb H_2_O_2_ redox homeostasis through transient catalase inactivation, especially in contexts associated with disturbed metal homeostasis. These findings partly explain some deleterious side effects resulting from the use of D-Penicillamine as a therapeutic agent.

## Experimental Procedures

### Materials

All chemicals were purchased from Sigma-Aldrich and used as-is. Bovine liver catalase and superoxide dismutase (SOD) from bovine erythrocytes were also purchased from Sigma-Aldrich. Hydrogen peroxide solution (9.79 M) TraceSELECT Ultra was purchased from Fluka. Nitrones used in this study were a kind gift from Dr. F. Peyrot (Université de Paris, France), with the exception of 5,5-dimethyl-1-pyrroline *N*-oxide (DMPO) that was purchased from Sigma-Aldrich.

### Enzymatic assays

CAT activity was measured on an Uvikon 941 spectrophotometer (Kontron) or a Cary 300 Scan (Varian) equipped with a temperature-controlled water bath (±0.1 °C) by following the disappearance of H_2_O_2_ at 240 nm over 30-60 s at 25 °C, as previously described (**22**). Briefly, experiments were carried out in a 3 ml quartz cuvette containing 11.7 mM H_2_O_2_ in 50 mM phosphate buffer, pH 7.4 ± 1 mM diethylene triamine pentaacetic acid (DTPA). The reaction was initiated by adding 0.3-0.6 pmoles of CAT. The effect of D-Pen on the activity of CAT was determined from a solution of CAT (30-60 nM) preincubated at 25 °C for 120 min as a function D-Pen concentration (0-5 mM in the presence of DTPA or 0-0.5 mM in the absence of DTPA) in 50 mM phosphate buffer, pH 7.4 ± 1 mM DTPA. Noteworthy, CAT contains a 2.124±0.386 molar excess of nonspecific iron metal ions, as previously determined (**22**). Then, 10 μl of the reaction mixture was added to the 3 mL cuvette containing the H_2_O_2_ solution to initiate the reaction. Similar experiments were realized in the presence of 0.1 M DMPO or 0.4 mM bicyclo[6.1.0]nonyne (BCN). The relative half inhibitory concentration (IC_50_) values were obtained by plotting the relative activity of CAT as a function of D-Pen concentrations, and by fitting the data with a four-parameter logistic equation: A=A_min_+(A_max_−A_min_)/(1+([D-Pen]/IC_50_)^n^), where A_max_ is the maximal activity of CAT (constrained at 100%), A_min_ is the minimum activity achieved at saturating concentration of D-Pen and n is the Hillslope that characterizes the slope of the curve at its midpoint. Each experiment was performed at least in triplicate.

### UV-visible Spectroscopy

UV-Visible spectra were recorded on an Uvikon 941 spectrophotometer (Kontron) equipped with a water bath (±0.1 °C). UV-visible spectral changes over time induced by D-Pen reactivity with catalase were recorded as previously described (**22**) in a cuvette containing catalase (3-5μM) in 50 mM phosphate buffer, 1 mM DTPA pH 7.4 at 25°C. The reaction was initiated by adding D-Pen (2 mM). The kinetic parameters associated with the reaction between catalase and D-Pen were obtained by plotting the relative absorbance at 404 nm (disappearance of native catalase) or 426 nm (appearance of compound II) as a function of time, and by fitting the data with a triple or double exponential function, respectively. Similar experiments were carried out in the presence of various nitrones such as 5-tert-butoxycarbonyl 5-methyl-1-pyrroline *N*-oxide (BMPO, 10 mM), 5-diethoxyphosphoryl-5-methyl-1-pyrroline *N*-oxide (DEPMPO, 50 mM), 5-diisopropoxyphosphoryl-5-methyl-1-pyrroline *N*-oxide (DIPPMPO, 50 mM) or DMPO (100 mM), increasing amount of SOD (0-200 U) or BCN (0.4 mM). The kinetics for compound II reduction to native catalase were determined in separate experiments by adding buffer, DMPO (100 mM), BCN (0.4 mM) or SOD (200 U) to a reaction mixture containing catalase (5 μM) that already reacted for 200-220 min with D-Pen (2 mM) in 50 mM phosphate buffer, 1mM DTPA pH 7.4 at 25°C. The relative absorbance at 404 nm was then plotted as a function of time and the data were fit with a sigmoidal function: A = A_min_ + (A_max_-A_min_)/(1+e^-(t-t0.5)/n^), where A_max_ is the absorbance of native catalase (constrained at 1), A_min_ is the absorbance of catalase after 220 min reactivity with D-Pen, *t*_0.5_ is the half-life of compound II, and n describes the steepness of the curve. The rate constant for compound II reduction *k*_obs_ were then derived from *t*_0.5_ using the linear relation: *t*_0.5_=ln2/*k*_obs_.

### Stopped-Flow UV-visible Spectroscopy

The inactivation kinetics with D-Pen were briefly analyzed on a sequential stopped-flow apparatus SFM-3 from Bio-Logic Science Instruments at ICMMO (Université Paris-Saclay, France). The following experiments were performed mainly to study the influence of SOD on compound I formation: catalase (4 μM final) and D-Pen (2 mM final) solutions were mixed together in the absence or presence of 200 U SOD (final) in 50 mM phosphate buffer, 1 mM DTPA pH 7.4. The kinetic data were recorded over 300 s or 620 s and analyzed using the BIO-KINE software application.

### HPLC-Mass spectrometry analysis

HPLC-MS (High Performance Liquid Chromatography coupled to Mass Spectrometry) spectra were recorded on a Thermo-Finnigan Surveyor equipped with a Satisfaction RP18AB C_18_ 3 μm (15×2 mm) (Cluseau) and coupled to an ESI LCQ Advantage (Thermo Fisher Scientific). The running mixture was subjected to filtering on a Microcon filter unit of 10 kDa (15 min, 4°C, 13,000 rpm) and the analyses were conducted on the flow-through. The HPLC separation was achieved at 0.2 mL/min using the following steps: 0-20 % B (0-15 min), 20 % B (15-25 min), 20-60 % B (25-30 min), 60 % B (30-50 min), 60-0 % B (50-51 min), 0 % B (51-60 min), with A = 0.1 % formic acid in water (or 10 mM ammonium acetate buffer, pH 4.6) and B = acetonitrile. Under these conditions, the retention time for D-Pen disulfide and the adduct of D-Pen-BCN were 7.5 min and 27.7 min, respectively.

### EPR spectroscopy

EPR measurements were performed on a Bruker Elexsys 500 EPR spectrometer operating at X-band (9.85 GHz) and equipped with a SHQ high-sensitivity cavity. Typical settings were: modulation frequency, 100 kHz; modulation amplitude 0.2 mT; receiver gain, 60 dB; time constant, 40.96 ms; conversion time, 41.04 ms; sweep width, 15 mT and sweep time, 42.02 s. EPR spectra were recorded sequentially during the whole reaction course at 21°C. Typical experiments were performed on a 50 μL solution containing 50 μM catalase, 2 mM D-Pen and nitrones at 50 mM (DEPMPO) or 200 mM (DMPO) in 50 mM phosphate buffer, 1 mM DTPA pH 7.4 at room temperature. Data acquisition and processing were performed using Bruker Xepr software. Appropriate control experiments (nitrone ± catalase, D-Pen or potassium ferricyanide) were performed.

## Results

### Catalase is inhibited by D-penicillamine

At first, we monitored CAT activity as a function of the concentration of D-penicillamine (D-Pen) in the presence or absence of the chelating agent diethylene triamine pentaacetic acid (DTPA). In the presence of DTPA, CAT is partially inactivated by D-Pen with a relative IC_50_ value of 191±26 μM (n=9±SD) (**Fig. 1A**) as well as a bimolecular rate constant of 21.5±2.7 M^-1^min^-1^ at 25°C (**Fig. S1A**). The partial inactivation of CAT (48.4±0.9 %) supports half-site reactivity, as previously observed with aminotriazole (**24**) or some biological thiols (**22**). As observed, D-Pen is a better inhibitor than the latter that display relative IC_50_ values (obtained in similar conditions) of 0.59±0.07 mM, 1.44±0.37 mM and 21.2±8.5 mM for cysteine (Cys), homocysteine (HCys) or glutathione (GSH), respectively (**22**).

**Figure 1.**
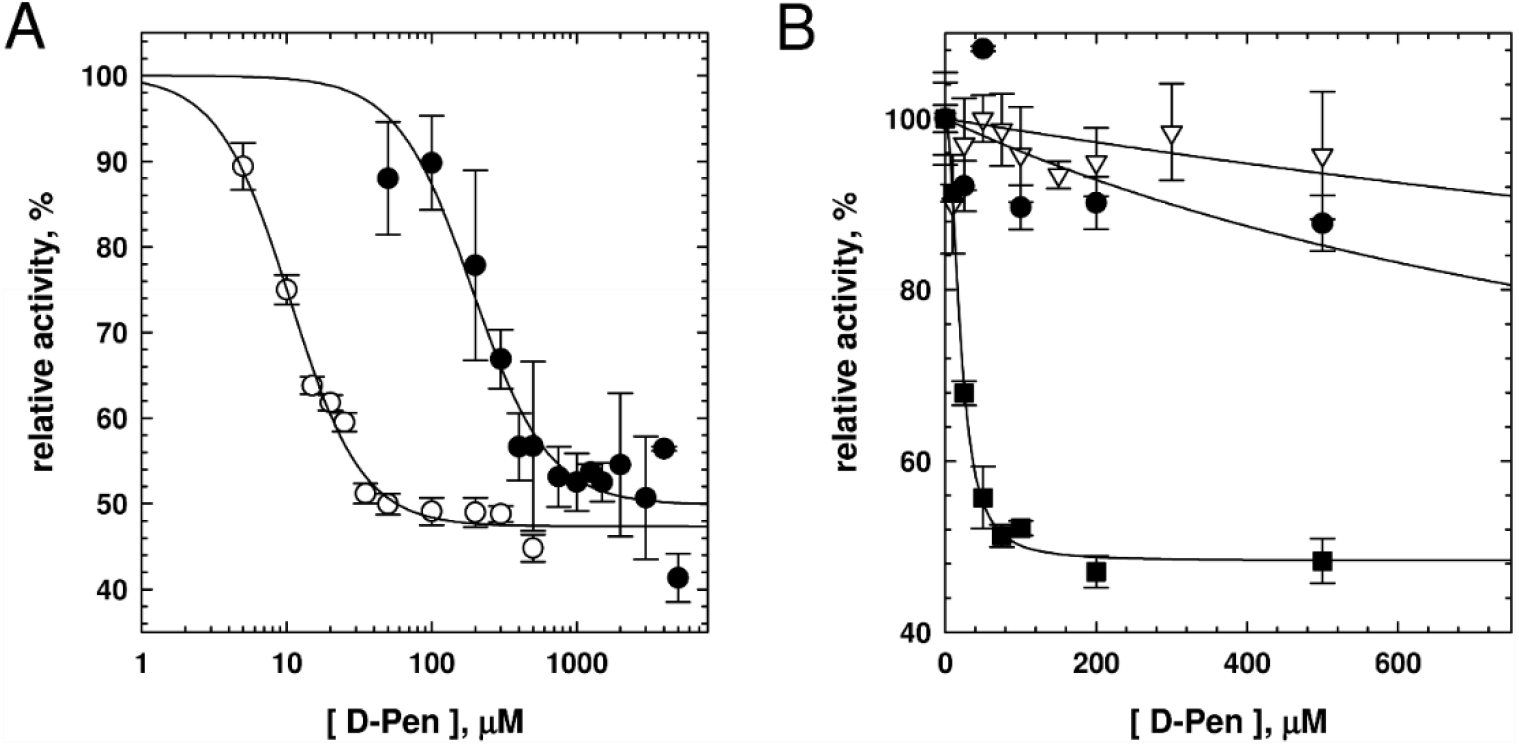
Inhibition of catalase bioactivity by D-penicillamine. (**A**) Experiments showing the dependency of CAT activity on the concentration of D-Pen in the presence (black symbols) or absence (white symbols) of 1 mM DTPA in 50 mM phosphate buffer at pH 7.4 and 25 °C. The plots were fitted with a four-parameter logistic equation and the relative IC_50_ values (n≥3±SD) are reported in the text. (**B**) Dependency of CAT activity as a function of D-Pen concentration in the absence of DTPA (black squares) and in the presence of 0.1 M DMPO (black circles) or 0.4 mM BCN (white triangles) in 50 mM phosphate buffer at pH 7.4 and 25 °C (n=3±SD).

In the absence of DTPA, the relative IC_50_ value drops to 10.6±0.5 μM (n=3±SD) (**Fig. 1A**). This strongly suggests that sulfhydryl compounds and redox-active transition metal ions (in here iron metal ions) may cooperate to mediate the inactivation of catalase bioactivity. To determine which species is responsible for the aforementioned decrease in the relative IC_50_ value, we monitored the activity of CAT as a function of the concentration of D-Pen in the presence of iron metal ions (64-128 nM), used to mimic hepatic non-heme iron overload, and various additives (**Fig. 1B**). D-Pen exhibits relative IC_50_ values greater than 1.25 mM in the presence of the nitrone spin-trap 5,5-dimethyl-1-Pyrroline-N-Oxide (DMPO) or the bioconjugating agent bicyclo[6.1.0]nonyne (BCN), suggesting that thiyl radicals and/or superoxide radical anions (O_2_^•-^) promote the inhibition of CAT activity (**Fig. 1B**). In addition, superoxide dismutase (SOD) accelerates the initial inhibition velocity of CAT by D-Pen (*v*_i_ = 0.71±0.11 min^-1^ or *v*_i_ = 1.10±0.15 min^-1^ (n=2±SD) in the absence or presence of SOD, respectively) (**Fig. S1B**), strongly advocating for hydrogen peroxide (H_2_O_2_) intervention during D-Pen-induced CAT inhibition. This observation is reminiscent of the inhibition of CAT bioactivity by biological thiols (**22**).

### D-penicillamine induces compound II formation

Next, we monitored spectral changes over time when CAT is allowed to react with D-Pen in the presence of DTPA (**Fig. 2A**). The UV-visible spectra recorded during the time course of the reaction clearly show the formation of several distinct species. The first transient species is formed within the minute, as observed by UV-visible spectroscopy (**Fig. 2A**) and confirmed by UV-visible stopped-flow spectroscopy (**Fig. S2**). This species exhibits distinct spectral features (α-band close to 660 nm) characteristic of compound I, the porphyrin radical-ferryl state of catalase [CAT-Fe(IV)=O]^• +^ (**25**). This intermediate is transformed into a second transient species typical of compound II, the one-electron oxidized ferryl state of catalase CAT-Fe(IV)=O (Soret band at 421, α-bands at 528 and 567 nm) (**22**). Noteworthy, our rapid kinetic experiments failed to detect the appearance of a band at 411 nm or 414 nm that are characteristic of the CAT-Fe(II) and CAT-Fe(II)-O_2_ species, respectively (**26**). In addition, we did not observe any other intermediate species than those already described.

**Figure 2.**
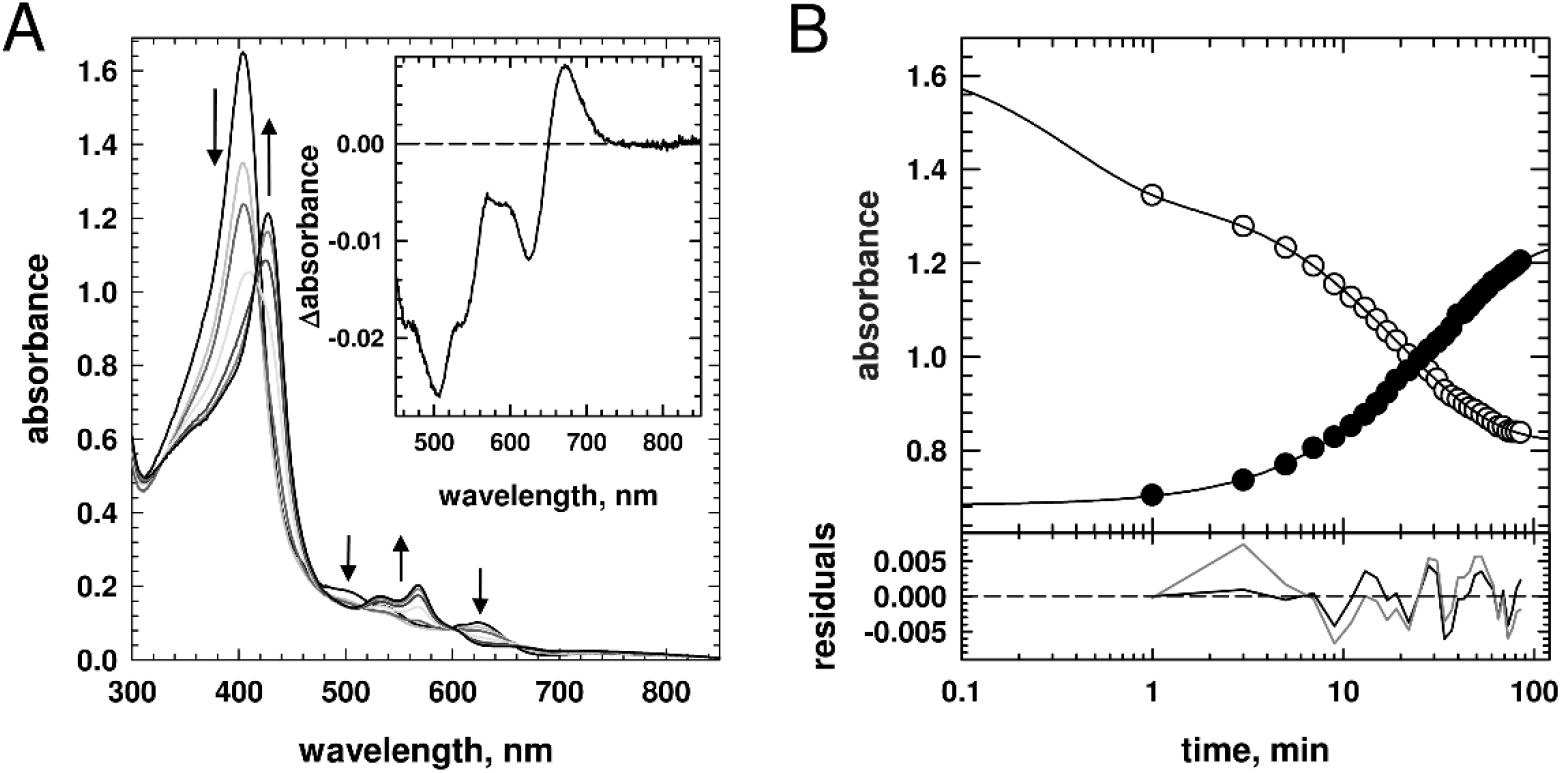
D-penicillamine induces compound II formation. (**A**) Representative experiment showing the UV-visible spectral changes recorded over time (0-89 min) when catalase (5 μM) reacts with D-Pen (2 mM) in 50 mM phosphate buffer, pH 7.4, 1 mM DTPA at 25 °C. Inset: Difference spectrum observed after catalase reacted for 1 min with D-Pen as in (A). The spectral peak around 660 nm is characteristic of compound I [CAT-Fe(IV)=O]^• +^. (**B**) Kinetics extracted from (A) at 404 nm (white symbols) and 426 nm (black symbols). The kinetic data at 404 nm were best fitted to a triple exponential function with *k*_obs1_ = 2.83±0.34 min^-1^, *k*_obs2_ = 0.0996±0.0274 min^-1^ and _*k*obs3_ = 0.0380±0.0073 min^-1^ as suggested by the residual plots below the graph. The dark and grey lines are the residual plots for a fit obtained with a triple or double exponential function, respectively. The kinetic data at 426 nm were best fitted to a double exponential function (dark trace) with *k*_obs1_ = 0.0613±0.0382 min^-1^, *k*_obs2_ = 0.0229±0.0105 min^-1^.

### D-penicillamine oxidation mediates the formation of compound II through two distinct mechanisms

Next, we focused on obtaining insights into the formation of compound II by performing kinetic analyses. Kinetic analyses of CAT reactivity with D-Pen show that native CAT-Fe(III) disappearance recorded at 404 nm is best fitted with a triple exponential function, while CAT-Fe(IV)=O appearance recorded at 426 nm can be best fitted with a double exponential function (**Fig. 2B**). In addition, both rates and relative amplitudes at 426 nm are close to those observed for the last two stages at 404 nm. These results thus strongly suggest that compound II, responsible for CAT inactivation, is generated through two apparent distinct routes respectively characterized by *k*_obs2_ = 0.0787±0.015 min^-1^ and *k*_obs3_ = 0.0443±0.0110 min^-1^ (n=7±SD) (**Table I**).

**Table I.**
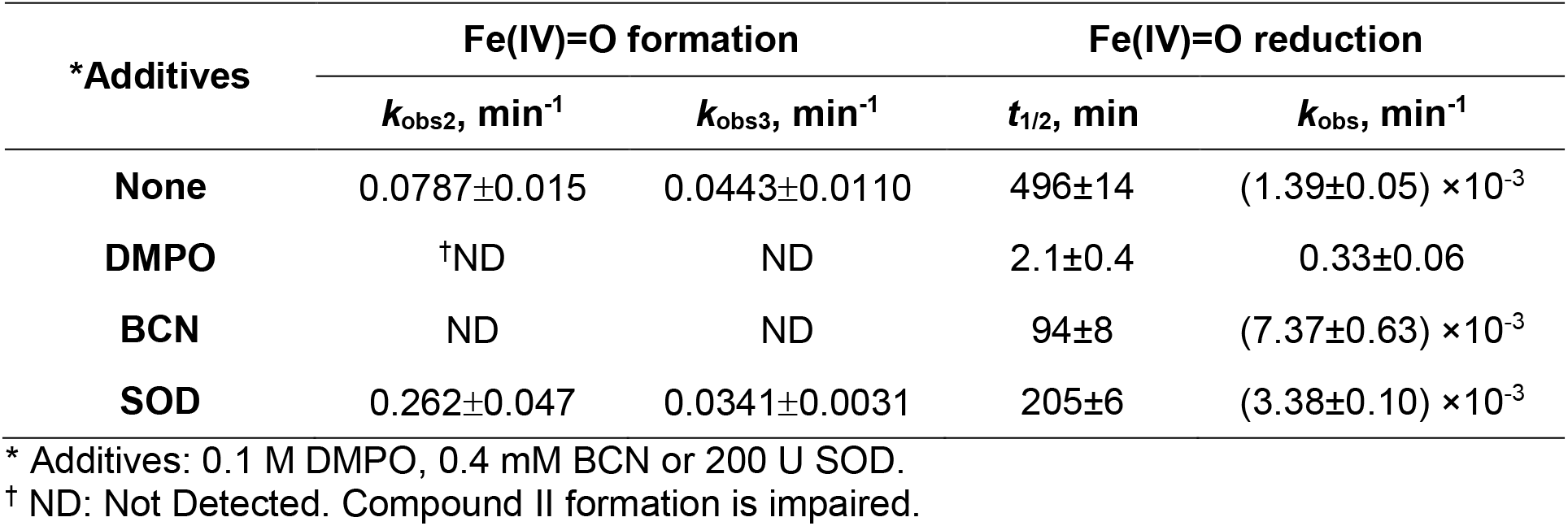
Rate constants for the reactivity of catalase with D-penicillamine at 25°C.

| *Additives | Fe(IV)=O formation |  | Fe(IV)=O reduction |  |
| --- | --- | --- | --- | --- |
| | $k_{\text{obs}2}, \text{ min}^{-1}$ | $k_{\text{obs}3}, \text{ min}^{-1}$ | $t_{1/2}, \text{ min}$ | $k_{\text{obs}}, \text{ min}^{-1}$ |
| **None** | $0.0787 \pm 0.015$ | $0.0443 \pm 0.0110$ | $496 \pm 14$ | $(1.39 \pm 0.05) \times 10^{-3}$ |
| **DMPO** | <sup>†</sup>ND | ND | $2.1 \pm 0.4$ | $0.33 \pm 0.06$ |
| **BCN** | ND | ND | $94 \pm 8$ | $(7.37 \pm 0.63) \times 10^{-3}$ |
| **SOD** | $0.262 \pm 0.047$ | $0.0341 \pm 0.0031$ | $205 \pm 6$ | $(3.38 \pm 0.10) \times 10^{-3}$ |

\* Additives: 0.1 M DMPO, 0.4 mM BCN or 200 U SOD.
<sup>†</sup> ND: Not Detected. Compound II formation is impaired.

Noteworthy, the preferential route depicted by *k*_obs3_ (ΔA_3_ = (1.54±0.42)×ΔA_2_, n=7±SD) exhibits a rate close to the ones registered for CAT-Fe(IV)=O generation with similar concentrations of biological thiols, thus suggesting that this pathway may be associated with compound II production through compound I reduction by superoxide radical anions (**22**). To better decipher the two distinct mechanisms responsible for compound II formation, we performed similar experiments to those described above in the presence of various additives, as detailed below.

### Intervention of D-penicillamine-derived thiyl radicals during compound II formation

The reaction between native catalase and D-penicillamine begins with the reduction of CAT-Fe(III) by D-Pen (**Fig. 3**). This process should initially occur through the formation of the Low Spin iron-complex CAT-Fe(III)-D-Pen, then the deprotonation of the bound D-Pen by the distal histidine followed by the rapid reduction of CAT-Fe(III) to CAT-Fe(II) along with a D-Pen derived thiyl radical generation. As observed by UV-visible spectroscopy (**Fig. 2A**), both aforementioned CAT intermediates are not detected during the time course of CAT-Fe(IV)=O formation. So we focused on demonstrating the production of D-Pen derived radicals during the generation of compound II. Accordingly, we first examined spectral changes over time during the reactivity of CAT and D-Pen with BCN, with subsequent analysis of the reaction mixture by HPLC-MS/MS, or with various nitrone spin trapping agents (DMPO, BMPO, DEPMPO or DIPPMPO) (**Table I, Fig. 4A-C and S3A**).

**Figure 3.**
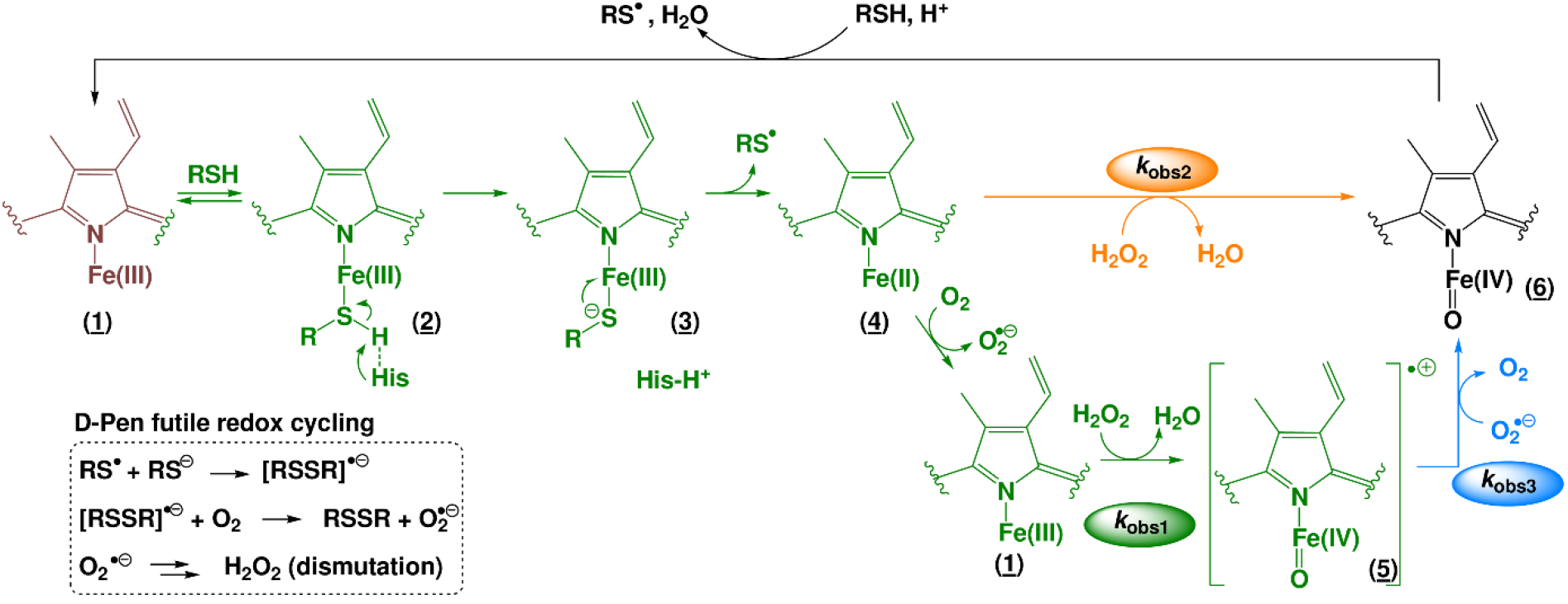
Mechanisms for the reversible formation of compound II induced by D-penicillamine. The reaction between native CAT-Fe(III) and D-Pen (RSH) starts with the reduction of CAT-Fe(III) by D-Pen (steps (<u>1</u>)-(<u>4</u>)) through the generation of the Low Spin iron-complex CAT-Fe(III)-RSH (<u>2</u>), the deprotonation of the bound D-Pen by the distal histidine to generate an unstable CAT-Fe(III)-RS-complex (<u>3</u>) that is rapidly converted to reduced CAT-Fe(II) (<u>4</u>) along with a D-Pen-derived thiyl radical RS^•^. The latter enters in a futile redox cycle described in the frame. At this point, CAT-Fe(II) (<u>4</u>) can be oxidized by molecular oxygen to CAT-Fe(III) (<u>1</u>), which then reacts with H_2_O_2_ to form compound I (<u>5</u>). Noteworthy, the rate *k*_obs1_ measured at 404 nm during CAT reactivity with D-Pen (Fig. 2) corresponds to compound I formation through steps (<u>1</u>)-(<u>5</u>). Compound I is subsequently reduced to compound II (<u>6</u>) by superoxide radical anions with a rate *k*_obs3_. Alternatively, compound II can be unexpectedly produced from the reaction between CAT-Fe(II) (<u>4</u>) and H_2_O_2_ at a rate *k*_obs2_ that exhibits sensitivity to superoxide dismutase (SOD). Last, compound II (<u>6</u>) slowly decays back to native CAT-Fe(III) (<u>1</u>) with a rate also sensitive to SOD.

**Figure 4.**
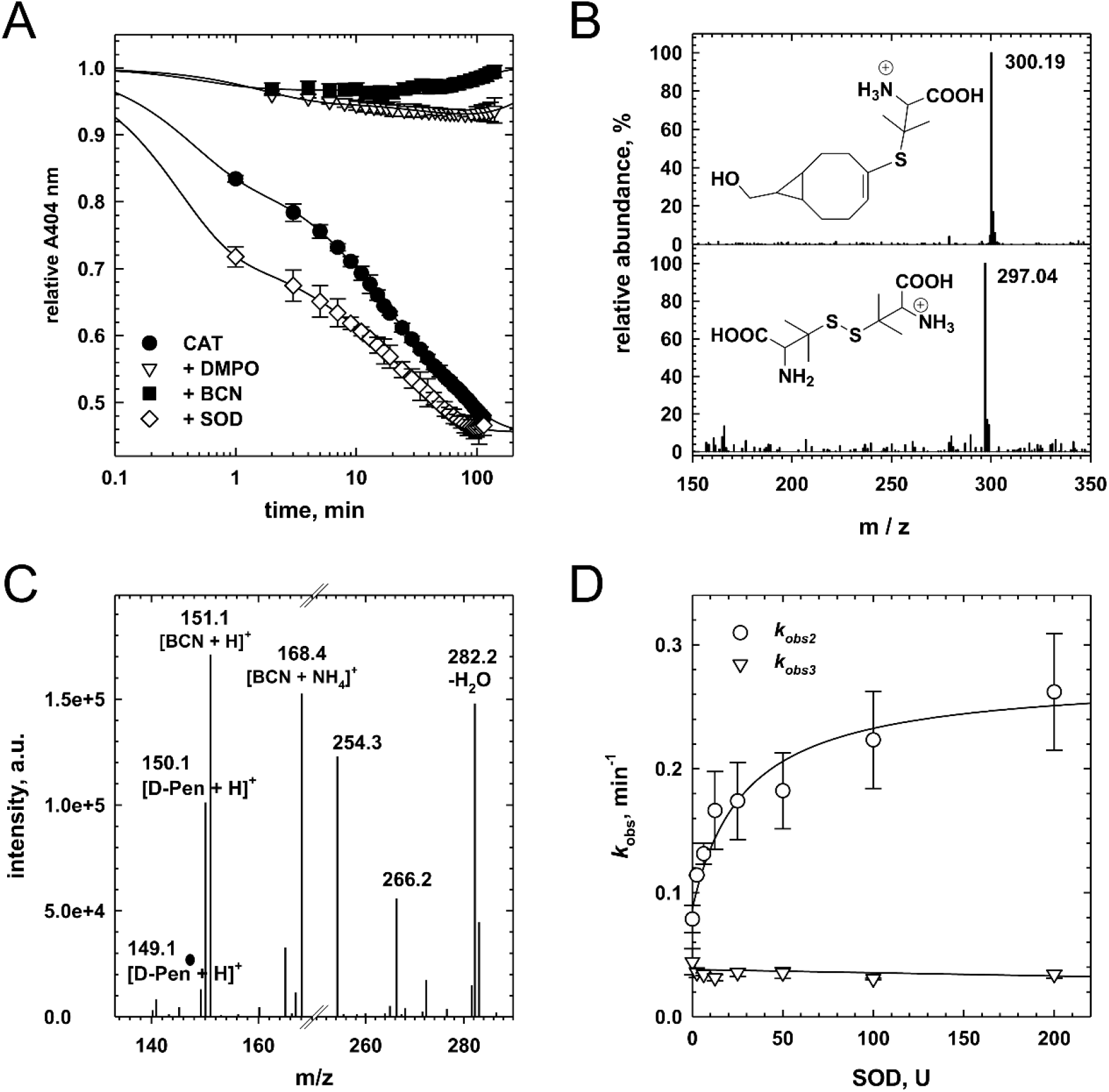
Sulfur and oxygen reactive species intervene during compound II formation. (**A**) Representative effect of various additives (100 mM DMPO, 0.4 mM BCN or 200 U SOD) on the kinetics of compound II formation during the reactivity of CAT (5 μM) with D-Pen (2 mM) in 50 mM phosphate buffer, 1 mM DTPA at pH 7.4 and 25 °C (n=2±SD). The kinetics for compound II formation in the absence of any additive (black circles) is shown for comparison. (**B**) Mass spectra (ESI+) of the BCN adduct (top) and the D-Pen disulfide (bottom) formed during the reactivity of catalase (4 μM) with D-Pen (2 mM) in the presence of 0.4 mM BCN in 50 mM phosphate buffer, 1 mM DTPA at pH 7.4 and 25 °C. (**C**) MS-MS spectrum (ESI+) of the molecular ion with a m/z = 300.2. The molecular ions at m/z = 266.2 and 254.3 result formally from the loss of H_2_O_2_ and a decarboxylation concomitant with the introduction of an unsaturation, respectively. The other main molecular ions are identified on the MS-MS spectrum. (**D**) Effect of increasing amount of superoxide dismutase (0-200 U) on the rates of compound II formation during the reaction of CAT (5 μM) with D-Pen (2 mM) in 50 mM phosphate buffer, 1 mM DTPA at pH 7.4 and 25°C. The rates *k*_obs2_ (white circles) and *k*_obs3_ (white triangles) have been extracted from the absorbance at 404 nm as described in Figure 2 (n≥2±SD).

The comparative kinetic studies realized in the presence of BCN or DMPO (**Table I, Fig. 4A**) or other nitrone spin trapping agents (**Fig. S3A**) clearly show that compound II formation is greatly impaired, thus strongly advocating for D-penicillaminyl radical intervention during CAT-Fe(IV)=O generation. The HPLC-MS/MS analysis of the reaction mixture obtained in the presence of BCN reveals the presence of a molecular ion at m/z=300.19 [M+H^+^] (**Fig. 4B**) that corresponds well to a BCN-D-Pen adduct (**Fig. 4C**). This adduct, not detected in the absence of the enzyme, results from the addition of D-Pen-derived thiyl radicals to BCN according to the thiol-yne reaction (**27**). Its presence in the reaction mixture thus clearly confirms the formation of D-penicillaminyl radicals through the oxidation of D-Pen by native CAT-Fe(III). Regrettably, even if the nitrone spin trapping agent DEPMPO strongly decelerates the rate of compound II formation (**Fig. S3A**) and the related reagent DMPO abolished CAT-Fe(IV)=O formation (**Fig. 4A**), we were unable to record any DMPO-or DEPMPO-D-Pen adducts by RT EPR spectroscopy, as observed with other biological thiols (**22**).

### Intervention of reactive oxygen species during compound II generation

Once formed, the D-Pen-derived thiyl radicals should enter into a futile redox cycle and partake into the production of reactive oxygen species (O_2_^•-^ and H_2_O_2_) along with D-Pen disulfide formation (**Fig. 3**). We thus analyzed by HPLC-MS the D-Pen derivatives formed during D-Pen reactivity with native CAT. Our analysis shows the presence of a molecular ion at m/z=297.04 [M+H^+^] that corresponds well to D-Pen disulfide (**Fig. 4B**). This observation confirms the intervention of D-penicillaminyl radicals into a futile redox cycle that will *in fine* produce O_2_^•-^ then H_2_O_2_ (**Fig. 3**), as observed during D-Pen-dependent copper-catalyzed H_2_O_2_ generation (**28**-**30**).

Then, the formation of compound II can occur *via* two different likely scenarios (**Fig. 3**). In the first pathway, compound II may be generated from the reaction between CAT-Fe(II) and H_2_O_2_, a mechanism observed during the non-enzymatic heme degradation by reactive oxygen species (**31**) and the heterolytical cleavage of H_2_O_2_ by ferrous myeloperoxidase (**32**) or ferrous lactoperoxidase (**33**). As noted above, the second one may be associated with compound II production *via* a similar mechanism than that observed with biological thiols, *i.e*. compound I reduction by superoxide radical anion (**22**). To discriminate between both pathways and correctly attribute the observed kinetic parameters to one mechanism or the other, we performed comparative kinetic studies of compound II formation in the presence of SOD, which should favor the formation of compound II through the first pathway.

At first, we examined the reactivity of CAT and D-Pen over time in the presence of 200 U SOD. The presence of SOD into the reaction mixture significantly accelerates compound I formation, as observed by UV-visible stopped flow spectroscopy (**Fig. S2A**), increases 3.3-fold the rate of compound II formation *k*_obs2_ and slightly decreases the rate of compound II generation *k*_obs3_ (**Table I, Fig. 4A**). Next, we performed similar comparative kinetic studies in the presence of increasing amount of SOD. As reported in **Fig. 4D**, while *k*_obs2_ exhibits hyperbolic dependence on SOD amount (*k*_min_ = 0.087±0.010 min^-1^, *k*_max_ = 0.275±0.028 min^-1^ and [SOD]_50_ = 28.9±10.1 U), *k*_obs3_ displays a linear dependence on it (slope = (−2.6±1.8)×10^−5^). Taken together, these results imply that *k*_obs2_ could be attributed to the formation of compound II through CAT-Fe(II) reactivity with H_2_O_2_, which is unusual and has never been demonstrated for CAT, and *k*_obs3_ could be associated with compound II formation through the reduction of compound I by O_2_^•-^, as observed with some biological thiols (**22**). Surprisingly, the amount of CAT-Fe(IV)=O generated through each respective pathway, represented here by the ratio ΔA2 over ΔA3, displays a bell-shaped curve as a function of SOD amount (**Fig. S2B**). Since the first pathway depends on H_2_O_2_ reactivity with CAT-Fe(II) and the second pathway relies on compound I reduction by O_2_^•-^, we would have expected to observe a linear or hyperbolic dependence of the ratio ΔA_2_/ ΔA_3_ as a function of SOD amount. Accordingly, this result implies that the proportion of compound II generated through each mechanism is finely adjusted by the amount of O_2_^•-^ *vs* H_2_O_2_ produced in the reaction mixture. It also suggests that the quantity and/or the flux of H_2_O_2_ modulates the selectivity of CAT-Fe(II) toward H_2_O_2_ or O_2_ oxidation.

### Compound II is unstable in the absence of thiyl radicals and reactive oxygen species

Next, we recorded the reactivity of catalase with D-penicillamine over a long period of time to determine if CAT-Fe(IV)=O is reduced back to native CAT-Fe(III), as observed with Cys or GSH, or is converted to another species, as detected with HCys (**22**). As reported in **Table I** and **Fig. S3B**, compound II slowly decays back to CAT-Fe(III) with a rate of (1.37±0.05) × 10^−3^ min^-1^. Noteworthy, the rate of decay to native catalase probably depends on a cycling between CAT-Fe(III) and compound II. If so, the more D-Pen-derived thiyl radicals and/or reactive oxygen species present in the reaction mixture, the longer will compound II be the dominating intermediate. To determine if that is the case we performed similar kinetic studies in the presence of various additives acting on the depletion of thiyl radicals or reactive oxygen species in the reaction mixture (**Table I, Fig. S3B**). The nitrone spin trap DMPO accelerates more than 230-fold the reduction of compound II to native CAT-Fe(III) with a *k*_obs_ of (330±2) 10^−3^ min^-1^, while BCN or SOD have a significant but less effect on its decay with *k*_obs_ of (7.42±0.72) × 10^−3^ min^-1^ and (3.32±0.11) × 10^−3^ min^-1^, respectively. These results thus confirm our assumptions.

## Discussion

We investigated here the reactivity of catalase toward D-penicillamine, a thiol that is potent in treating Wilson’s disease, rheumatoid arthritis and alcohol dependence. The results of our studies offer a clear basis for understanding the mechanism whereby D-penicillamine reversibly inhibits catalase, hence possibly affecting various cellular processes through disturbance of H_2_O_2_ levels. Thus, our data indicate that complex electron transfer reactions take place between catalase and/or non-heme iron, D-penicillamine and oxygen, promoting the generation of CAT-Fe(IV)=O, a temporarily inactive state of the enzyme, as well as reactive oxygen species and thiyl radical intermediates.

Numerous studies demonstrated that CAT bioactivity is either inhibited through the formation of 6-coordinate Low Spin (*e.g*. with cyanide) or 5-coordinate High Spin (*e.g*. with formate) heme-iron-complexes, the production of an unclear type I inactive catalase (*e.g*. with 2-mercaptoethanol), the generation of sulfheme (*e.g*. with homocysteine) or compound II (*e.g*. with cysteine or glutathione) (**22**,**34**-**37**). As reported in here, our data clearly indicate that CAT bioactivity is inhibited by D-Pen through the generation of compound II, as observed with Cys and GSH (**22**). However, D-Pen is a better inhibitor than biological thiols in term of half maximal inhibitory concentration IC_50_. This may be due to D-Pen higher relative hydrophobicity in comparison to the aforementioned biological thiols, as shown by their respective partition coefficient ClogP (Chemdraw 19.1, Perkin Elmer) of −3.293, −3.678, −8.589 or −2.585 for Cys, HCys, GSH and D-Pen, respectively. As a result, D-Pen could have better access to catalase active site. In addition, the lower p*K*_a_ of the thiol group in D-Pen compared to the other aforementioned thiols could also account for these results. Accordingly, the relative reactivity of D-Pen and biologically important thiols with O_2_^•-^ or H_2_O_2_ is inversely related to their p*K*_a_ (**38**). Noteworthy, the relative IC_50_ values exhibited by D-Pen, in particular the one displayed in the presence of iron metal ions used to mimic non heme iron overload, fall within the range of D-penicillamine concentration observed in plasma of patients receiving the drug (**30**).

Usually, at low levels of H_2_O_2_ and in the presence of one-electron donors (for example biological thiols, phenols, aromatic amines, NO or O_2_^•-^), compound I may go through a one-electron reduction towards the temporarily inactive state of the enzyme compound II, which converts back to native catalase over a second one-electron reduction step (**22**,**36**,**39**). Alternatively, compound II could be transformed into compound III, an oxyferrous state (CAT-Fe(II)-O_2_) of the enzyme that could go back to native CAT-Fe(III) along with O_2_^•-^ release. As expected, the former scenario is also followed in the presence of D-Pen, without the detectable presence of compound III. However, based on experiments carried out in the presence of SOD that accelerates (hyperbolic dependence, **Fig. 4D**) and increases (bell-shape curve, **Fig. S2B**) the formation of compound II, we also demonstrated that a surprising involvement of CAT-Fe(II) reactivity with H_2_O_2_ promotes compound II formation and competes with the classical pathway.

Previous investigations showed that the ferrous form of catalase is normally inaccessible by conventional reduction techniques of native catalase (**36**) and requires the three-electron reduction of compound I, for example by azide that promotes azydil radical formation and NO-ferrocatalase generation (**40**,**41**). Nevertheless, the present study underlines again the unique ability of thiols to reduce CAT-Fe(III) to CAT-Fe(II) (**22**). Also, contrary to some heme-peroxidase systems known to generate compounds II by the two-electron oxidation of their ferrous form by H_2_O_2_ (**32**,**33**), the ferrous form CAT-Fe(II) obtained by reduction of native catalase with biological thiols (Cys, HCys or GSH) rapidly decays back to native catalase by reacting with molecular oxygen rather than yielding compound II (**22**). Therefore, D-Pen induces a new type of reactivity for ferrous catalase, which was initially proposed to explain the mechanism of decomposition of hydrogen peroxide by catalase (**42**) but was quickly dismissed (**43**). This observation may be explained by the effect of substituents on the stability of sulfur-centered radicals (**44**,**45**) resulting from the reaction between the various thiols and native catalase. Accordingly, D-penicillamine-derived thiyl radicals could exhibit a gain in stability compared to thiyl radicals derived from biological thiols that could promote the participation of D-penicillaminyl radicals in the futile redox cycle described in **Fig. 3** and therefore enhance the production of superoxide. Additionally, Winterbourn *et al* showed that D-Pen produces more hydrogen peroxide upon reaction with superoxide than other biologically important thiols (**38**). The creation of a greater flux and/or amount of O_2_^•-^ then H_2_O_2_ in the presence of D-penicillamine could thus explain its surprising capacity to participate in the formation of compound II by a new mechanistic avenue, the two-electron oxidation of ferrous catalase. Thus, it appears that the reactivity of catalase with biologically relevant thiols is strongly tuned by their structure, in which slight changes result in diverse reactivities, as illustrated in **Fig. 5**.

**Figure 5.**
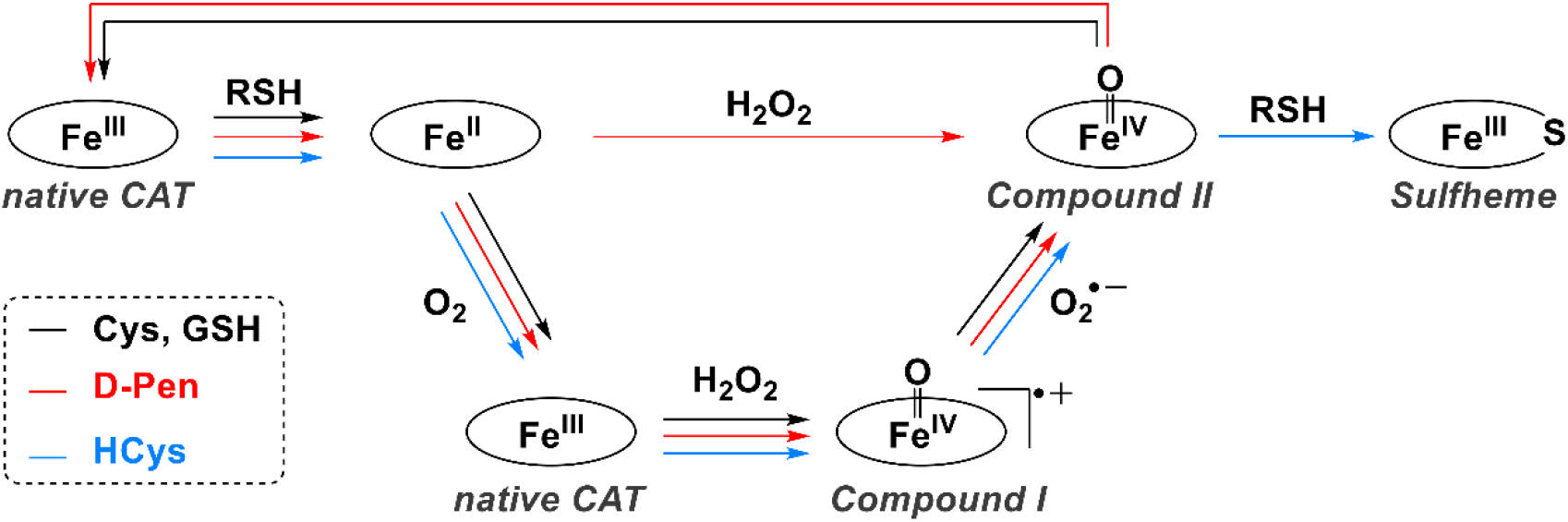
Reactivity of various biologically relevant thiols toward catalase.

Due to its chemical properties, D-penicillamine is used as a therapeutic copper chelating agent in the treatment of Wilson’s disease (**3**), as an immunosuppressor in the treatment of rheumatoid arthritis (**4**), and as a sequestering agent of acetaldehyde in the treatment of alcoholism (**9**). Unfortunately, adverse effects are fairly frequent with the use of D-penicillamine (**10**-**16**), and our study reveals a possible molecular basis for this Janus-faced behavior. Thus, we confirm the pro-oxidant capacity of D-penicillamine that leads to hydrogen peroxide generation by reaction with metal ions, which recently favored its repurposing in anticancer therapy (**28**,**29**) and may facilitate its action on the acute inflammatory response during treatment of RA (**30**). In addition, we show that the transient but recurring catalase inactivation initiated by D-Pen could in the long term perturb H_2_O_2_ redox homeostasis and signaling, based on the recent evidence that peroxisomes and catalase are central in the H_2_O_2_ signaling network (**23**,**46**). Last, a prolonged treatment with D-Pen could induce a chronic inflammatory and oxidative state through chronic disruption of H_2_O_2_ redox homeostasis. This phenomenon could explain the long-term presentations reported in patients treated with D-Pen, that is hepatic iron accumulation and a buildup of iron in the brain, which may be a risk factor for developing neurodegenerative diseases (**14**,**47**-**51**). Thus, iron uptake and cellular accumulation are favored by various H_2_O_2_-dependent mechanisms under pathological conditions, including H_2_O_2_-dependent up-regulation of transferrin receptor 1 (**52**), stimulation of hepcidin production and ferroportin internalization and degradation (**53**,**54**), and alteration of ceruloplasmin and its overall ferroxidase activity (**55**-**57**).

## Acknowledgment

We are indebted to Dr Fabienne Peyrot (Université de Paris, France) for precious assistance with EPR experiments, to Assia Hessani (now at Sorbonne Université, France) for mass-spectrometry analysis, and to Dr Jean-Noël Rebilly (Université Paris-Saclay, France) for useful support with UV-visible stopped-flow experiments.

