## Supplementary figures for "Molecular Basis for the Interaction of Catalase with D-Penicillamine : Rationalization of some of its Deleterious Effects"

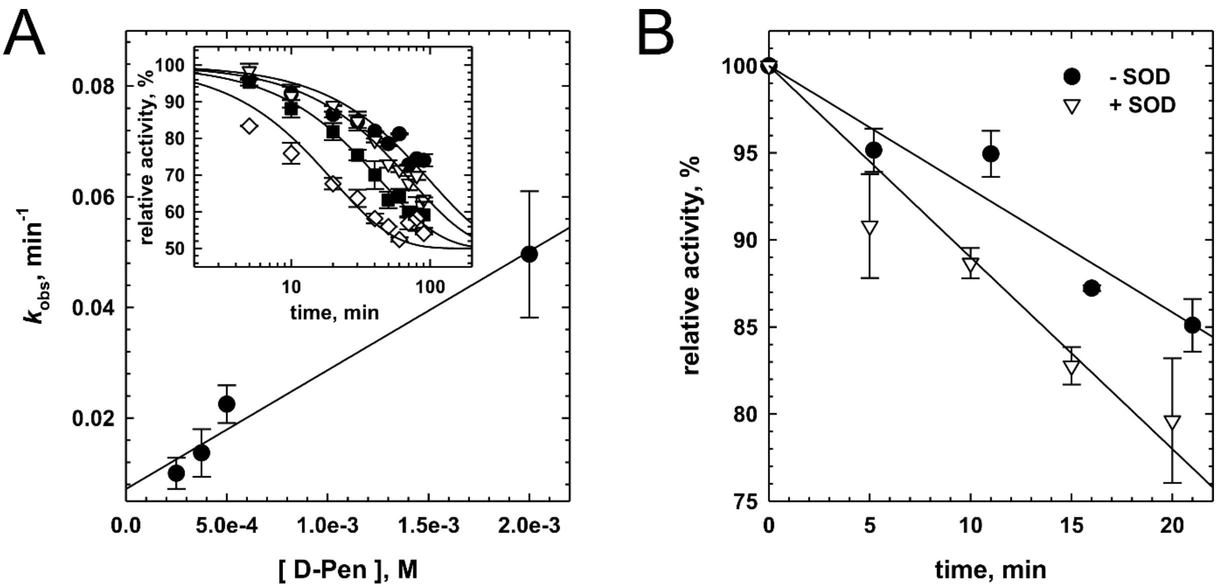


**Figure S1 - Catalase inhibition by D-penicillamine.** (**A**) Dependence of the inhibition rate of CAT bioactivity on the concentration of D-Penicillamine in 50 mM phosphate buffer, 1 mM DTPA at pH 7.4 and 25 °C. Inset: Time-dependent inhibition of CAT bioactivity by various amount of D-Penicillamine (0.25 (●), 0.375 (▽), 0.5 (■) and 2 (◇) mM) in 50 mM phosphate buffer, 1 mM DTPA at pH 7.4 and 25 °C. (**B**) Experiments showing the time-dependent inhibition of CAT activity by D-Pen (0.25 mM) in the presence (open symbols) or absence (dark symbols) of 100 U SOD in 50 mM phosphate buffer at pH 7.4 and 25 °C. The plots were fitted with a linear equation and the initial velocity values (n=2±SD) are reported in the text.

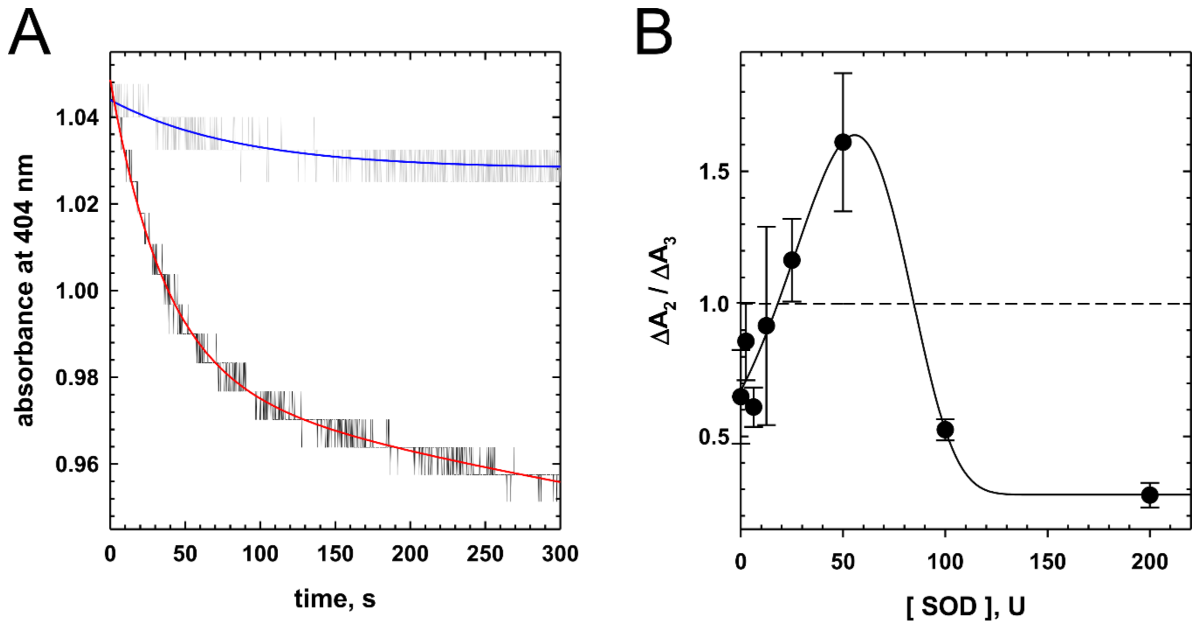


**Figure S2 - Effect of superoxide dismutase on D-penicillamine-induced compound I and compound II generation.** (**A**) Representative kinetic traces recorded at 404 nm during the reactivity of CAT (4 μM) with D-Pen (2 mM) in the presence (dark trace) or absence (grey trace) of SOD (200 U) in 50 mM phosphate buffer, 1 mM DTPA at pH 7.4 and 20°C. The kinetic data were best fitted to a double (in the presence of SOD - red trace; $k_{obs1}$ = 1.79±0.15 min$^{-1}$) or single (in the absence of SOD - blue trace; $k_{obs}$ = 0.74±0.08 min$^{-1}$) exponential function. (**B**) Effect of increasing amount of superoxide dismutase (0-200 U) on the amount of CAT-Fe(IV)=O generated through the pathway characterized by ΔA2 (formation through the reactivity of CAT-Fe(II) with $H_2O_2$) or ΔA3 (reduction of compound I by $O_2^{\bullet -}$). The data $\Delta A_2$ and $\Delta A_3$ have been obtained from the experiments represented in **Fig. 3D**.

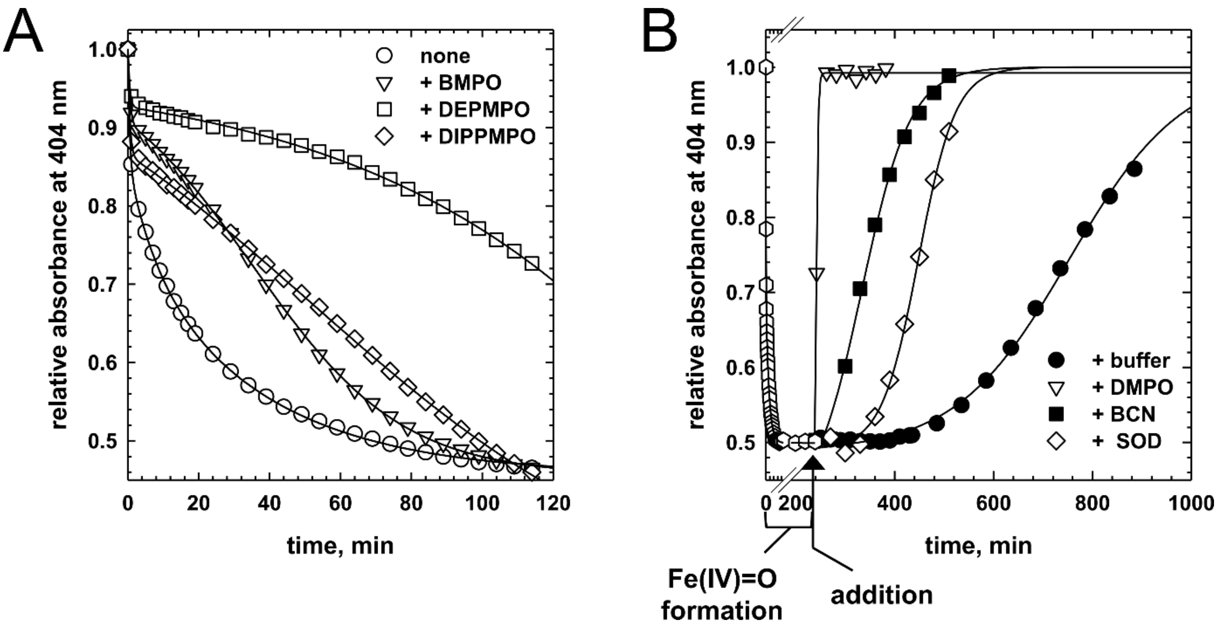


**Figure S3 - Representative effect of various additives on the kinetics of compound II formation or reduction.** (**A**) Representative effect of various nitrones (10 mM BMPO, 50 mM DEPMPO or 50 mM DIPPMPO) on the kinetics of compound II formation during the reactivity of CAT (4 μM) with D-Pen (2 mM). The kinetics for compound II formation in the absence of any additive (white circles) is shown for comparison. (**B**) Representative effect of various compounds (100 mM DMPO, 0.4 mM BCN or 200 U SOD) on the kinetics of compound II reduction after CAT (5 μM) reacted with D-Pen (2 mM) for 240 min (white hexagonal symbols). The kinetics for compound II reduction in the absence of any additive (black circles) is shown for comparison. The kinetic data were fit with a four parameters sigmoidal equation to extract the half-life ($t_{1/2}$) of compound II after the addition of the various additives. All the experiments were performed in 50 mM phosphate buffer, 1 mM DTPA at pH 7.4 and 25°C.
